# Photoinduced DNA Lesions in Dormant Bacteria. The Peculiar Route Leading to Spore Photoproduct Unraveled by Multiscale Molecular Dynamics

**DOI:** 10.1101/2020.04.28.065516

**Authors:** Antonio Francés-Monerris, Cécilia Hognon, Thierry Douki, Antonio Monari

## Abstract

Some bacterial species enter a dormant state in the form of spores to resist to unfavorable external conditions. Spores are resistant to a wide series of stress agents, including UV radiation, and can last for tens to hundreds of years. Due to the suspension of biological functions such as DNA repair, they accumulate DNA damage upon exposure to UV radiation. Differently from active organisms, the most common DNA photoproduct in spores are not cyclobutane pyrimidine dimers, but rather the so-called spore photoproduct. This non-canonical photochemistry results from the dry state of DNA and the binding to small acid soluble proteins that drastically modify the structure and photoreactivity of the nucleic acid. In this contribution, we use multiscale molecular dynamics simulations including extended classical molecular dynamics and QM/MM biased dynamics to elucidate the coupling of electronic and structural factors leading to this photochemical outcome. In particular, we rationalize the well-described impact of the peculiar DNA environment found in spores on the favored formation of the spore photoproduct, given the small free energy barrier found for this path. Meanwhile, the specific organization of spore DNA precludes the photochemical path leading to cyclobutane pyrimidine dimers formation.

**TOC GRAPHICS:** 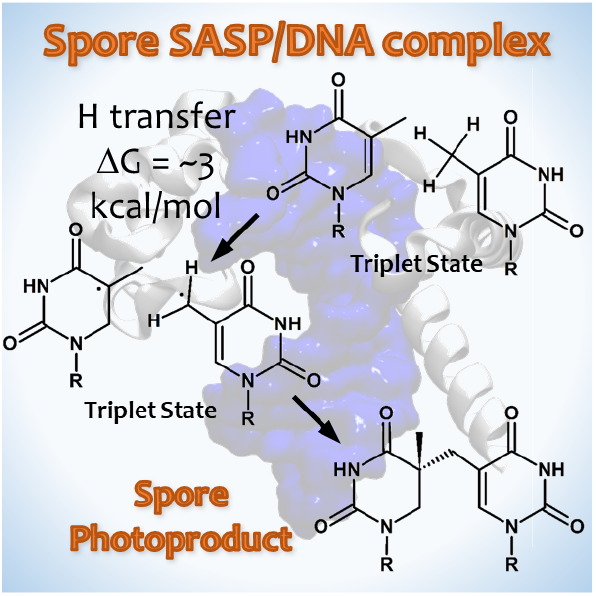

---

Spores represent a dormant form characteristic of some bacteria, that is used as a defense mechanism to cope with difficult or prohibitive environmental conditions such as shortage in nutrients or heat, and hence allow the organism survival.^1–5^ The spore state, that is characterized by the presence of additional walls within the cell and extensive dehydration, can last for tens or even hundreds of years until the environmental conditions become sufficiently favorable to trigger germination,^6^ which reinstates all physiological processes of the living organism. Spores are also characterized by an extreme resistance to many external stress agents, such as chemicals, ionizing radiations and UV light. Understanding how spores resist to intense UV fluxes raises some fascinating basic questions that may for instance be related to the emergence of life. In addition, survival of spores is a key issue in space exploration with the risk of cross-planetary contamination on spacecraft.^7,8^ Last, because of their outstanding resistance, spores represent an evident issue in all decontamination processes and in particular those using UVC, and consequently constitute a public health threat.^9,10^

**Scheme 1).**
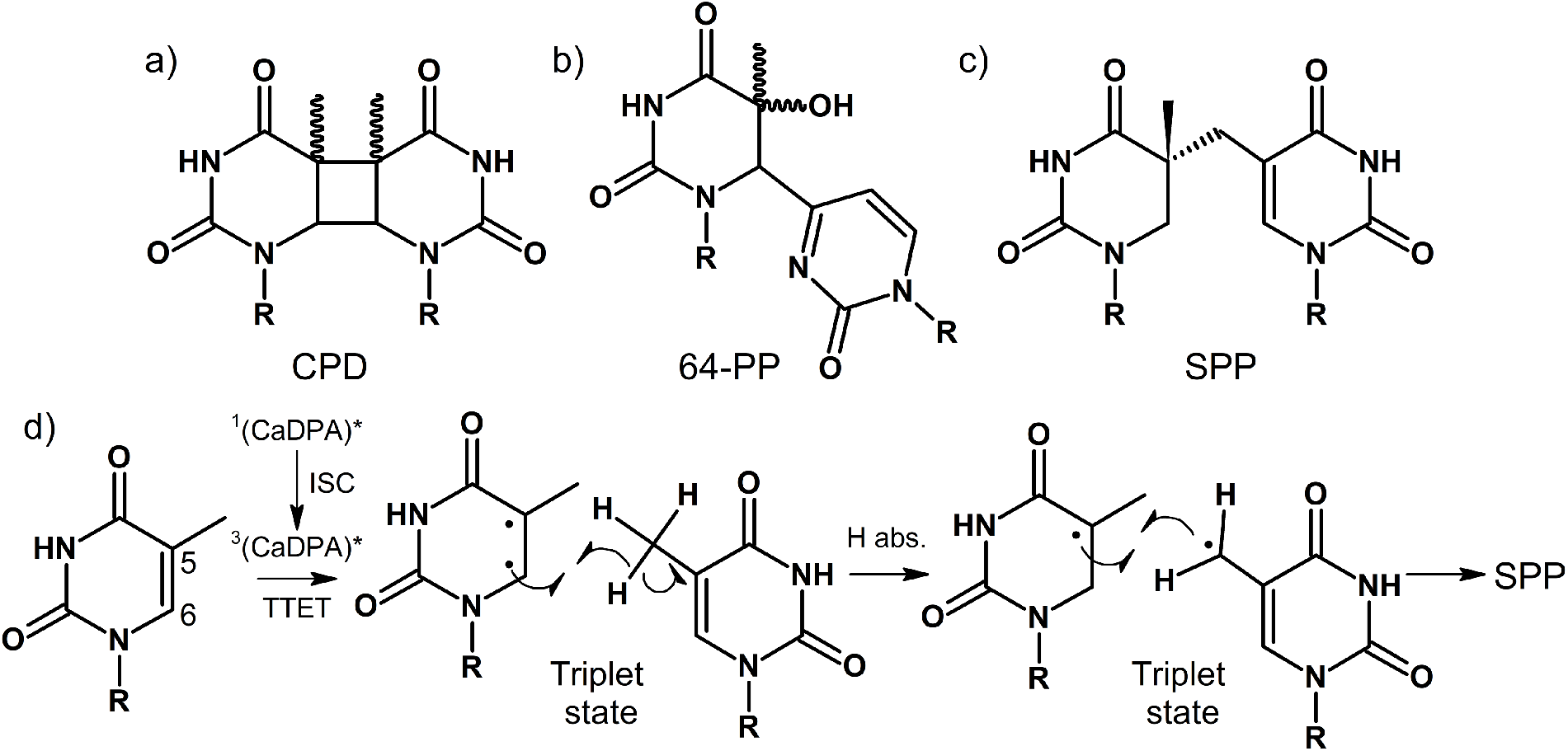
Structures of a) cyclobutane pyrimidine dimer (CPD), b) 64-photoproduct (64-PP), and c) spore photoproduct (SPP). Panel d) shows the stepwise molecular mechanism for the formation of SPP (see text).^14–16^

Due the prolonged exposure to solar radiation and the suspension of the normal cellular cycles, and hence of all DNA repair machinery, spores may be seen as a hotspot for the accumulation of photoinduced DNA lesions. Indeed, one of the critical steps of germination is precisely the activation of a fast and low-energy consuming DNA repair pathway, the very specific radical-SAM spore photoproduct lyase, to reinstate genome stability.^11–13^

While in active prokaryotes and eukaryotes organisms the main photoinduced DNA lesions are constituted by cyclobutane pyrimidine dimers (CPD) and in a lesser extent pyrimidine (6-4) pyrimidone photoproducts (64-PP), the almost exclusive lesion in spores, the so-called spore photoproduct (SPP),^16^ present a totally different chemical structure (Scheme 1) characterized by the intrastrand bridging of the methyl group of the 3’ thymine with the 5-carbon of the nearby 5’ thymine.^17^ SPP was first discovered in the sixties in UV irradiated spores^18^ and may be responsible of the death of the spore,^19,20^ but it is also related to its resistance thanks to the peculiar DNA repair mechanism that is activated upon spore germination, and specifically tailored to SPP removal.^13,21^

The fact that spores trigger new pathways for the formation of specific DNA lesions, that are almost never observed in other biological systems, calls for the elucidation of the specific role played by the organization of the DNA macromolecular environment in tuning the photophysical and photochemical outcomes.^1,22,23^ Indeed, and in addition to the very low water content, two peculiarities of the spores’ biological environment need to be highlighted. First, the center of the spore hosting the genetic material is saturated with a high quantity of pyridine-2,6-dicarboxylic acid chelated with Ca^2+^ ions (CaDPA). CaDPA presence has been associated with important functions assuring spore resistance, stability and germination,^6,24,25^ while it also exerts a protective role towards DNA avoiding its desiccation and deterioration.^26^ However, CaDPA may also act as a photosensitizer,^27^ populating its triplet state manifold through a facile intersystem crossing to further transfer the energy to DNA by means of triplet-triplet energy transfer, hence triggering DNA photoreactivity. Furthermore, the genomic DNA is enrolled around and protected by specific proteins, the small acid soluble proteins (SASP),^27–30^ that induce a considerable structural deformation as compared to solvated DNA. Notably, these factors are extremely specific of the spore environment and upon germination CaDPA is rapidly released from DNA while SASPs are readily degraded.^3,25,31^

The chemical mechanism of SPP formation has been the object of a number of experimental studies, involving both model systems and solid-state DNA oligomers.^27,32–34^ It has been proposed that, right after the population of the thymine triplet manifold, a hydrogen is transferred from the methyl group of the 3’ thymine to the C6 site of the 5’ thymine (see Scheme 1d). Subsequently, the singlet ground state is recovered yielding the new carbon-carbon bond and the corresponding SPP. Apart from the DNA photosensitization, the rate-determining step of the overall process is clearly the hydrogen transfer process, that in turn should also dictate the specificity of the SPP formation.

In this contribution, we aim at clarifying and identifying all the structural and electronic factors leading to the specific and almost absolute predominance of the SPP formation in spores. To this aim, we have resorted to a multiscale modeling including classical molecular dynamics (MD) simulations and hybrid quantum mechanics/molecular mechanics (QM/MM) protocols. In particular, we have simulated the behavior of a DNA decamer complex with SASP (Figure 1, pdb 2z3x)^28^ and compared with the behavior of the same oligonucleotide extracted from the protein complex and solvated without the protein (hereafter referred to as naked DNA) or initially constructed in ideal B-form (B-DNA).

**Figure 1).**
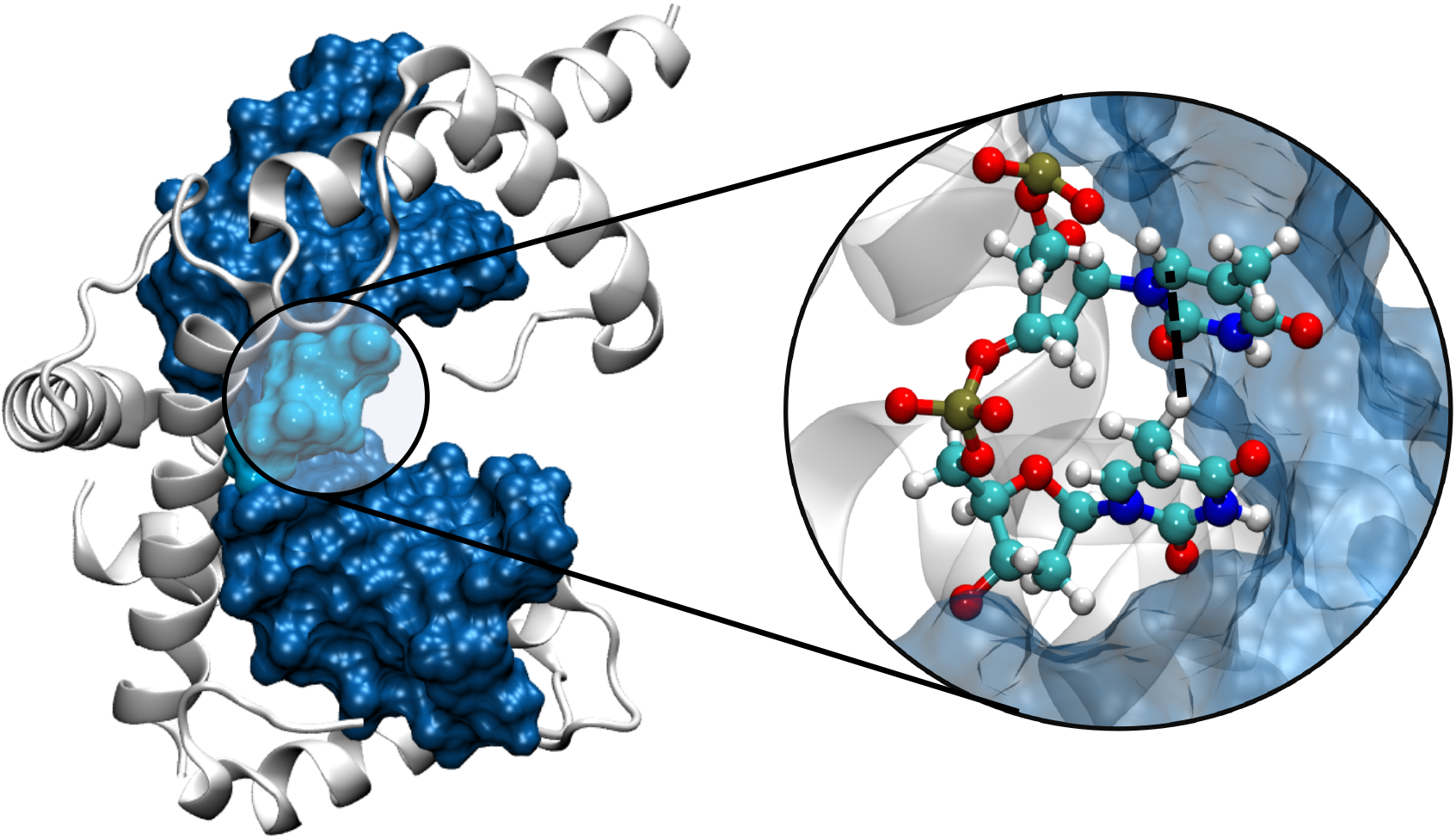
Representative snapshot of the MD simulations showing the complex between the DNA oligomer and SASP. The position of the two adjacent thymine moieties are shown in the zoom in ball and stick representation. The black dashed line highlights the reaction coordinate.

Two nucleobases from the original crystal structure were replaced by two thymine nucleobases at the center of the oligomer to allow for a potentially reactive site. Classical MD simulations were performed using the NAMD code^35^ and visualized via VMD,^36^ DNA and the protein were modeled using amberf99 force field including the bsc1 corrections,^37–39^ while water was described with the TIP3P force field.^40^ DNA structural parameters have been quantified using the Curves+ code.^41^ Subsequently, the hydrogen transfer reaction was studied both at full QM level in a model system composed of two p-stacked thymine nucleobases in the gas phase and by means of QM/MM simulations starting from snapshots extracted from the classical MD trajectory. Gas-phase calculations were performed with the GAUSSIAN 09 software^42^ whereas the QM/MM simulations were performed with the Amber 16 program^43^ interfaced with the ORCA 4.0 code.^44–46^ The potential of mean force (PMF) corresponding to the free energy profiles of the H transfers were computed through umbrella sampling techniques following a protocol used in previous works.^47,48^ The QM partition was composed of the two reactive nucleobases described at the density functional theory (DFT), making use of the ωB97-XD functional^49^ and the 6-31G basis set, given the very good accuracy provided by this level of theory (see Electronic Supporting Information, ESI). The reaction coordinate was set as the distance between the transferable hydrogen and the accepting carbon (see Figure 1 and ESI). Extensive details on the computational protocols and strategies are presented in ESI.

The MD simulation confirms the formation of a stable aggregate between DNA and SASP. The complex is quite rigid as witnessed by the very small value of the root mean square deviation (RMSD) exhibited by both the DNA and the protein fragment (Figure 2).

**Figure 2).**
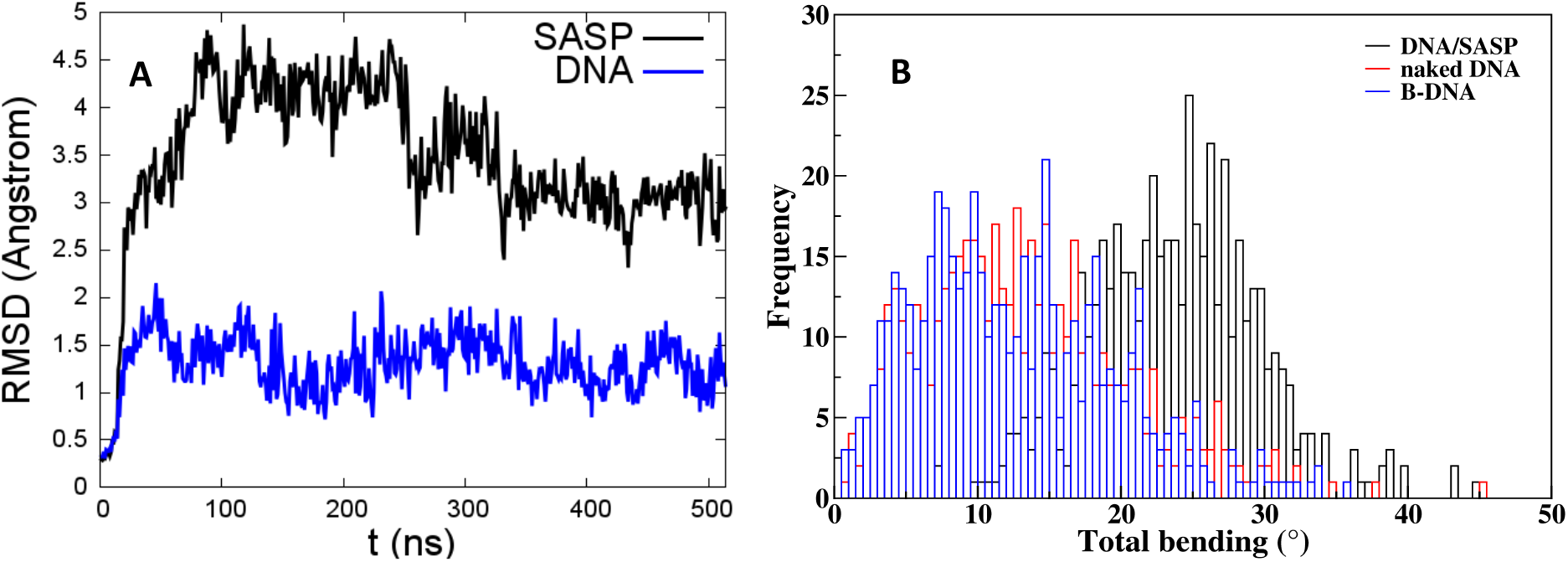
Time series of the RMSD over the MD simulation for the complexed SASP and DNA oligomer (A), and the distribution of the DNA bending angle over the MD simulation for the DNA/SASP complex, B-DNA and naked DNA (B).

In agreement with what is observed for other DNA compacting proteins,^50^ the main driving force for the formation of the complex is due to favorable electrostatic interactions between the negatively charged DNA backbone and the positive protein residues, mainly arginine and lysine. Interestingly, and differently from the characteristic of more globular compaction agents, such as histones or histone-like proteins, SASP presents positively charged a-helices that accommodate in the DNA minor groove providing additional stabilization. Most probably, such evolution is a consequence of the extremely water-poor environment of the spores, requiring special neutralization and protection of the DNA groove.

Despite the stability of the DNA-SASP complex, the snapshot reported in Figure 1 evidences that SASP induces a strong structural deformation on the DNA strand, strongly bending the nucleic acid and inducing a pronounced curvature. Indeed, the DNA form in SASP is closer to A-DNA, coherently with different experimental observations.^2^ It is important to note that this important bending is exerted on a rather short DNA oligomer, contrarily to the case of DNA compaction around histone proteins, in which the curvature spreads on a much longer fragment. Figure 2B compares the distribution of the bending for the DNA/SASP complex with the one for B-DNA and naked DNA. While the DNA/SASP complex clearly presents a peak at around 25°, both solvated oligomers, coherently with the behavior of canonical DNA, are less bent and their distribution peaks at around 10°. Furthermore, the distribution of the bending angles is much larger for naked and B-DNA than for the DNA/SASP complex, highlighting once more the global rigidity of the latter structure. SASP is also known to induce a twist of the helical turn of the DNA, that is nicely reproduced by our MD simulation, and that is claimed to produce a structure that lies in between the B- and A-form. The evolution of other structural parameters is also reported in ESI and shows, in addition to the bending, a global torsion of the DNA helix, evidenced in the backbone and the roll parameters, that also increases the interbase separation thus reducing the electronic coupling between the nucleobases and hampering canonical CPD mechanisms. The increase of the minor groove depth and the corresponding decrease of its width, due to the presence of the SASP, is also noticeable.

The distribution of the distances between the transferable hydrogen, i.e. one of the three hydrogens belonging to the methyl group of the 3’ thymine, and the accepting C6 carbon, largely overlaps that of the SASP/DNA complex and both solvated oligonucleotides, peaking at around 2.9-3.0 Å as reported in Figure 3A. In sharp contrast, the distribution of the distance between the center of mass of each thymine carbon-carbon double bond differs considerably between the three systems. Most notably, the average inter-thymine distance is larger for the SASP/DNA complex (~3.5 Å) than for the B-DNA (~2.5 Å). Moreover, the distribution is much broader in the case of the solvated oligonucleotide, once again pointing to the important increased rigidity of the DNA protein complex. Interestingly, the naked DNA does not recover the characteristic of the B-DNA strand presenting a peak in the distribution of the thymine-thymine distance comparable to the one of SASP/DNA although with a larger dispersion of the values.

**Figure 3).**
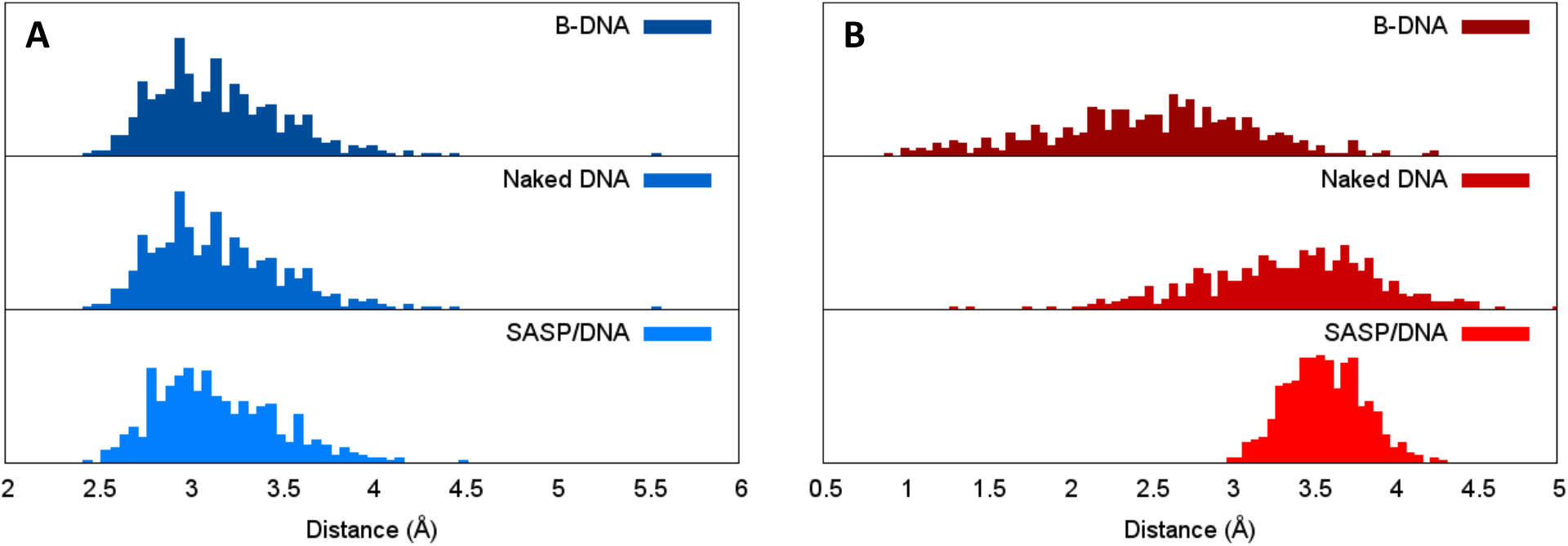
Distribution of the distances between the transferable hydrogen and the accepting carbon (A) and between the centers of mass of the thymine carbon-carbon double bond (B) for SASP/DNA, naked DNA, and B-DNA. See ESI for a better definition of the coordinates.

The increase of the thymine-thymine distance and the overall rigidity of the SASP/DNA structure can also be correlated with the absence of CPD lesions in this system. This is not surprising since large thymine-thymine distances destabilize excimer states delocalized over several stacked nucleobases, which represent the electronic states driving the CPD production in canonical DNA.^51–55^ Conversely, short thymine-thymine stacking distances stabilize excimer states delocalized over several nucleobases. Hence, the complex equilibrium between structural elements and electronic factors, and their coupling with the inhomogeneous environment, is crucial to tune the yield of SPP vs. CPD.

The reaction profiles for the SPP formation in the triplet state, calculated in a model system composed of two π-stacked nucleobases in a conformation similar to the one assumed in the B-DNA environment are reported in the SI. The agreement between the different levels of theory used is remarkable, all profiles show an exergonic reaction for the hydrogen transfer with the photoproduct lying about 10 kcal/mol lower in energy than the reactant. However, the process takes place with a significantly high energy barrier of about 15-20 kcal/mol, indicating a relatively slow reaction in absence of environment. The analysis of the transition state nature and the associated intrinsic reaction coordinate profile confirms that the reaction coordinate can be safely described as the carbon-hydrogen distance.

Figure 4 shows the QM/MM free energy profiles along the hydrogen transfer reaction coordinate obtained for the SASP/DNA complex and the solvated B-DNA oligomer. In the case of the SASP/DNA complex, it becomes apparent that while the reaction is still exergonic (3.2 kcal/mol), the free energy barrier is now strongly reduced and amounts to only 2.9 kcal/mol. This fact may be traced back to the influence of the environmental effects captured by the QM/MM simulations, such as the structural deformations shown in Figures 2 and 3. Hence, it is evident that the reaction is strongly favored in spore biological conditions as compared to the gas-phase and canonical B-DNA situation, justifying the observed predominance of the SPP in irradiated spores.

**Figure 4).**
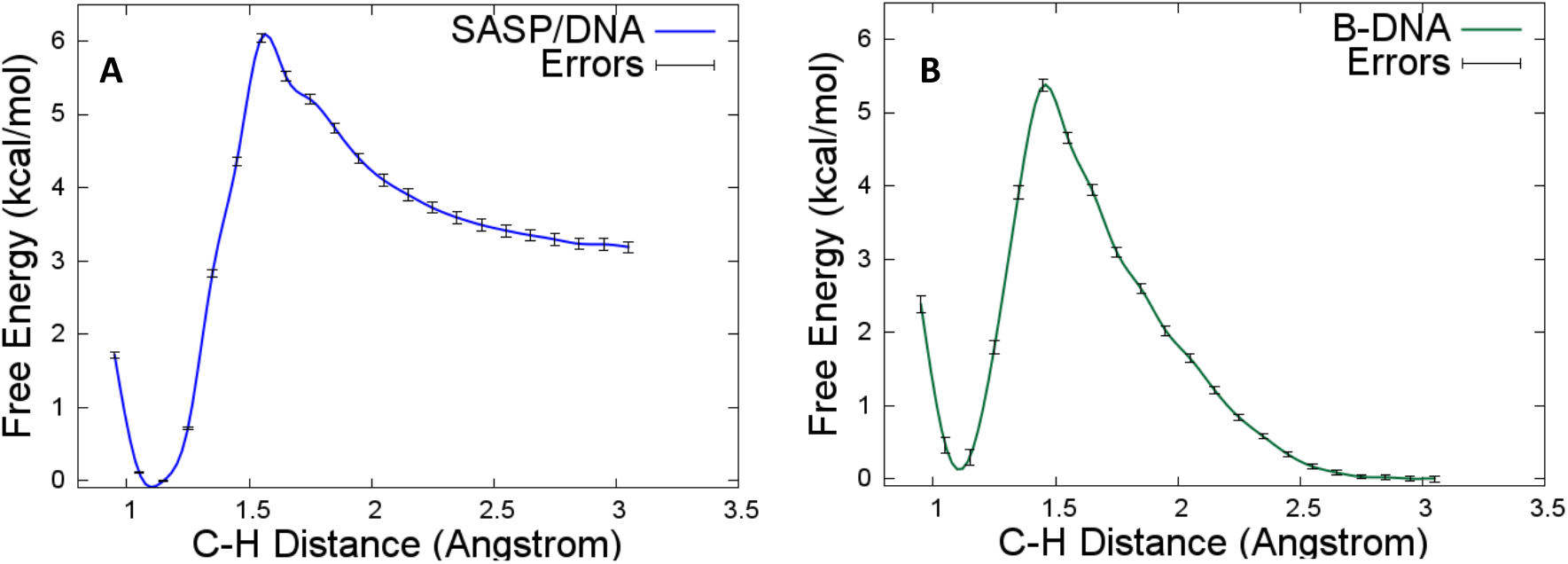
QM/MM free energy profile for the hydrogen transfer reaction obtained for the SASP/DNA complex (A) and the solvated DNA strand (B). The PMF was constructed from the ensemble of C-H bond distances using the weighted histogram analysis method (WHAM) developed by the Grossfield lab.^56^

Representative snapshots extracted from the most relevant areas of the DNA/SASP PMF, i.e. reactant, product, and transition state are reported in Figure 5 for the DNA/SASP complex and in ESI for B-DNA. No particular interactions with the protein environment or with the DNA backbone can be evidenced from the analysis of the simulation windows. As expected, the hydrogen loss induces the planarization of the thymine methyl radical. Note that the allyl radical could also further evolve via a tautomerization with the C-C double bond opening a further competitive channel, however the study of this process is out of the scope of the present contribution. Importantly, one can notice that the stacking of the nucleobases is conserved, hence providing the optimal conditions for the further formation of the C-C bond necessary to yield SPP, as displayed in Scheme 1d. Apart for the planarization of the methyl radical, no evident structural deformations are found neither in the transition state nor in the products, further justifying the observed small free energy barrier also in terms of a negligible reorganization energy penalty.

**Figure 5).**
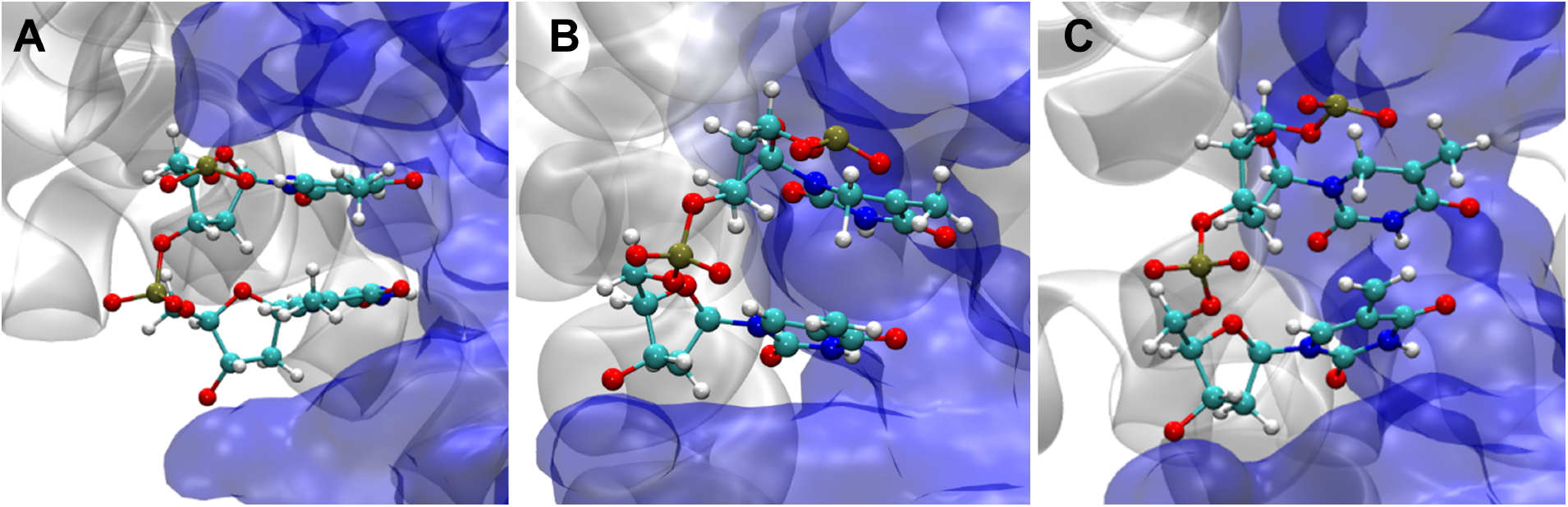
Representative snapshots extracted along the reaction coordinate for the reactant (A), transition state (B), and product (C) regions. DNA is shown in blue whereas SASP is displayed in white.

Important differences in the PMF obtained for the solvated B-DNA strand should be highlighted. Indeed, the thermodynamic driving force observed for the DNA/SASP complex is now absent and the reactant is greatly stabilized. As a matter of fact, the reaction is now slightly endergonic (0.3 kcal/mol), in stark contrast to the DNA/SASP reaction (see Figure 4). Furthermore, the free-energy barrier is higher and amounts to 5.4 kcal/mol. Both factors clearly evidence that the SPP formation is favored by the specific SASP environment as compared to the canonical B-DNA disposition.

The results obtained from our multiscale protocol rationalize the reasons behind the almost exclusive formation of SPP in spore. First, the presence in the vicinity of DNA of an effective photosensitizer, CaDPA,^27^ allows the population of the thymine triplet state manifold. Second, we have shown that the rate limiting step of the SPP formation, i.e. the hydrogen transfer, is characterized by only a very small free energy barrier (less than 3 kcal/mol), mostly as a result of the compaction of DNA around SASP. The condensation around SASP induces structural modifications leading to larger distances between the thymine C5=C6 double bonds and increasing the rigidity of the DNA double strand, thereby hampering the CPD formation. This conclusion is also reinforced by the free energy barrier leading to SPP, which is much lower in the case of the DNA/SASP complex than in the case of B-DNA. A global scenario can be sketched from all these data. In solvated DNA, the CPD pathways, on the triplet and singlet manifolds, are accessible and favorable, while the SPP formation is slightly penalized, hence CPD are the dominant lesions. Conversely, in spore, the DNA condensation around SASP penalizes the CPD pathway, while the SPP still requires only a very small, and biologically compatible, energy barrier thus becoming the most probable photoproduct.

Our results shed light on the still open question of the molecular and biological mechanisms of DNA lesions in spores and, more in general, constitute an example of the subtle equilibrium between electronic, environmental, and structural effects dictating the prevalence of a certain class of DNA lesion. In the following, we plan to study the whole SPP formation process and to quantitatively establish the free energy penalty for the CPD formation in the spore environment.

## ASSOCIATED CONTENT

### Supporting Information

Full computational details, IRC profiles, DNA parameters of the classical MD runs, histograms of the C-H distances for the simulation windows, snapshots of the QM/MM simulations for the B-DNA system.

## ACKNOWLEDGMENT

Support from the Université de Lorraine and CNRS is gratefully acknowledged. Calculations have been performed on the LPCT local computing cluster and on the regional computing center ExpLor in the framework of the project “Dancing under the light”. A. F.-M. is grateful to Generalitat Valenciana and the European Social Fund (APOSTD/2019/149) and the Ministerio de Ciencia e Innovación (project CTQ2017-87054-C2-2-P) for financial support.

