## Supplementary Information for "Photoinduced DNA Lesions in Dormant Bacteria. The Peculiar Route Leading to Spore Photoproduct Unraveled by Multiscale Molecular Dynamics"

AUTHOR INFORMATION

**Corresponding Authors**

*

### Full computational details

#### Molecular dynamics simulations

The crystal structure of the SASP/DNA complex was extracted from the Protein Data Bank pdb code 2Z3X. The nucleobases at the 16 and 17 positions, cytosine in the original structure, have been *in silico* mutated to thymine. The mutated DNA oligomer extracted from the original complex was also studied (naked DNA). In parallel, the same 10-bp oligomer having an ideal B-DNA structure (B-DNA) was built with the amber nucleic acids builder (nab) utility for comparison purposes.

The three systems (SASP/DNA, naked DNA, and B-DNA) were solvated in a cubic box containing ~10,000 TIP3P water molecules corresponding to an initial size of 79*77*61 Å$^3$. To assure electroneutrality, $K^+$ ions were added to balance the global negative charge.

All the simulations have been performed using the NAMD 2.13 code[1] with amber parmf99[2] force field including bsc1[3] correction for nucleic acid. The following protocol was consistently used for all the MD simulations: 1,000 steps of energy minimization to remove possible bad contacts, followed by a 6 ns equilibration progressively removing positional constraints on protein and DNA in the isobaric-isothermal ensemble (300K, 1 atm), and finally a production run. The latter was run for a total of 200 ns to allow the correct sampling of the conformational space of the systems and the assessment of their stability. To increase the performance, we used the Hydrogen Mass Repartition (HMR) method consisting in scaling all the hydrogen masses from 1.008 to 3.024 a.u., thus allowing the use of a 4.0 fs time step to integrate the Newton equations of motion.

Representative snapshots were analyzed and rendered using VMD,[4] while Curves+ program[5] was used to post process the MD simulations and obtain the DNA structural parameters.

#### Gas-phase calculations

The model composed by two thymine molecules has been extracted from a snapshot of the classical MD simulation described above. Reactant, transition state, and product have been optimized in the gas phase by means of the unrestricted DFT ansatz, making use of the long-range and dispersion-corrected ωB97-XD functional[6] in conjunction with the 6-31G, 6-31G*, and 6-311++G(d,p) basis sets, as implemented in the GAUSSIAN 09 software.[7] Analysis of the normal modes of the optimized geometries verifies the true nature of the transition states, showing only one imaginary frequency in all cases corresponding to the H transfer between the methyl group of one thymine to the C6 site of the adjacent thymine molecule. The reaction profiles have been calculated at the corresponding levels of theory by means of the intrinsic reaction coordinate (IRC) method to ensure the connectivity between reactant and product. In addition, the 6-31G IRC profile have been recomputed by means of the ORCA 4.0 program[8] without reoptimizing the structures, i.e. through single-point calculations. Results are shown in Figure S1. All profiles are coincident, indicating a negligible dependence of this reaction on the basis set and validating the use of the most efficient 6-31G basis set for the QM/MM biased simulations.

#### QM/MM biased simulations

The Amber 16 program[9] interfaced with the ORCA 4.0 program[8] was used to run all QM/MM simulations. This means that the Amber 16 program performed all nuclear displacements using the forces of the QM partition computed by ORCA 4.0. The QM-MM cutoff was set to 9 Å. The QM subspace comprised the two thymine nucleobases, whereas the rest of DNA strand, SASP protein, water molecules and $K^+$ atoms were treated with MM force fields as described in the MD details section. The QM level of theory was ωB97-XD/6-31G according to the benchmark described above and shown in Figure S1. We used the link atom procedure to saturate the valence of the system by adding hydrogens in the N1 atom of the thymine molecules and the C1' atoms of the sugar moieties.

The simulation windows used to sample the C-H reaction coordinate ranged from 1.00 to 3.00 Å and were separated by 0.1 Å, making a total of 21 windows for each potential of mean force (PMF). An extra window at 1.55 Å was included in the PMF of the SASP/DNA system to ensure the adequate overlap of all windows (see Figure S2). The C-H bond distances were kept by applying a force constant of 1200 kcal/mol. The windows were prepared as follows. First, 400 minimization steps were run at 1.50, 2.00, 2.50, and 3.00 Å starting from the snapshot extracted from the classical MD simulation. Later, the system was thermalized during 8 ps up to a temperature of 300 K in the NVT ensemble, followed by 2 ps of additional simulation. The last coordinates were subsequently used as initial structures (seeds) for the rest of windows repeating the aforementioned minimization/equilibration protocol. Finally, production runs of simulation times between 20 and 25 fs in the NPT ensemble were performed for each simulation window.

The PMF was constructed from the ensemble of C-H bond distances using the weighted histogram analysis method (WHAM) developed by the Grossfield lab.[10]

(2) Cornell, W. D.; Cieplak, P.; Bayly, C. I.; Gould, I. R.; Merz, K. M.; Ferguson, D. M.; Spellmeyer, D. C.; Fox, T.; Caldwell, J. W.; Kollman, P. A. A Second Generation Force Field for the Simulation of Proteins, Nucleic Acids,

and Organic Molecules. *J. Am. Chem. Soc.* **1995**, *117* (19), 5179–5197. https://doi.org/10.1021/ja00124a002.

(10) Grossfield, A. WHAM: An Implementation of the Weighted Histogram Analysis Method, Version 2.0.9. http://membrane.urmc.rochester.edu/content/wham/.

#### Gas-phase IRC profiles

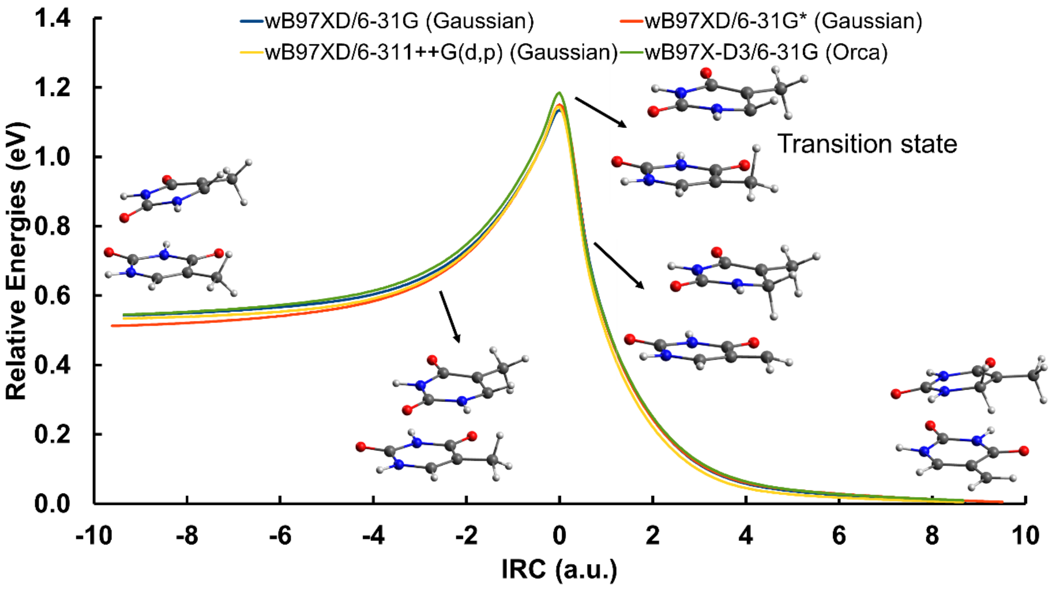

**Figure S1.** IRC profiles for the H transfer reaction in the triplet state for the model system in the gas phase.

### Histograms of the QM/MM windows

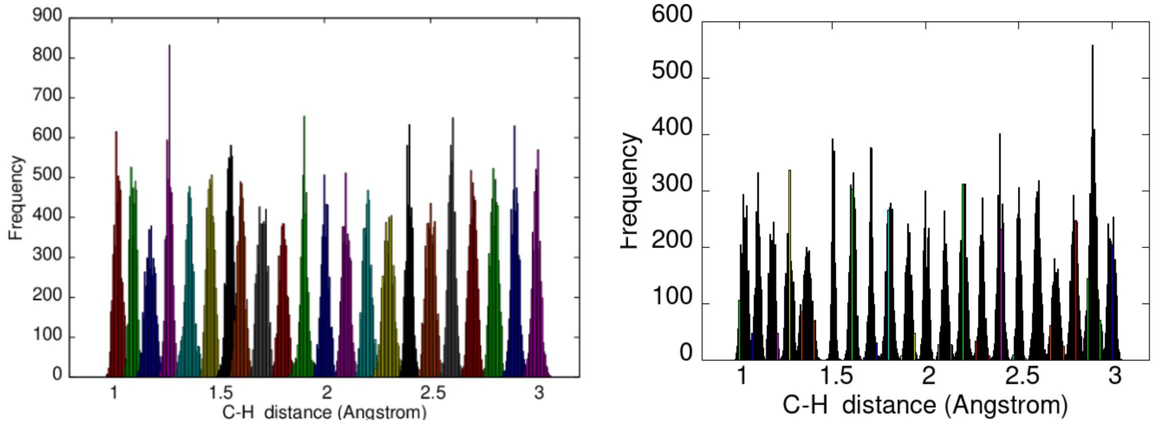

**Figure S2.** Histograms of the QM/MM windows belonging to the SASP/DNA (left) and B-DNA (right) PMFs.

### DNA structural parameters

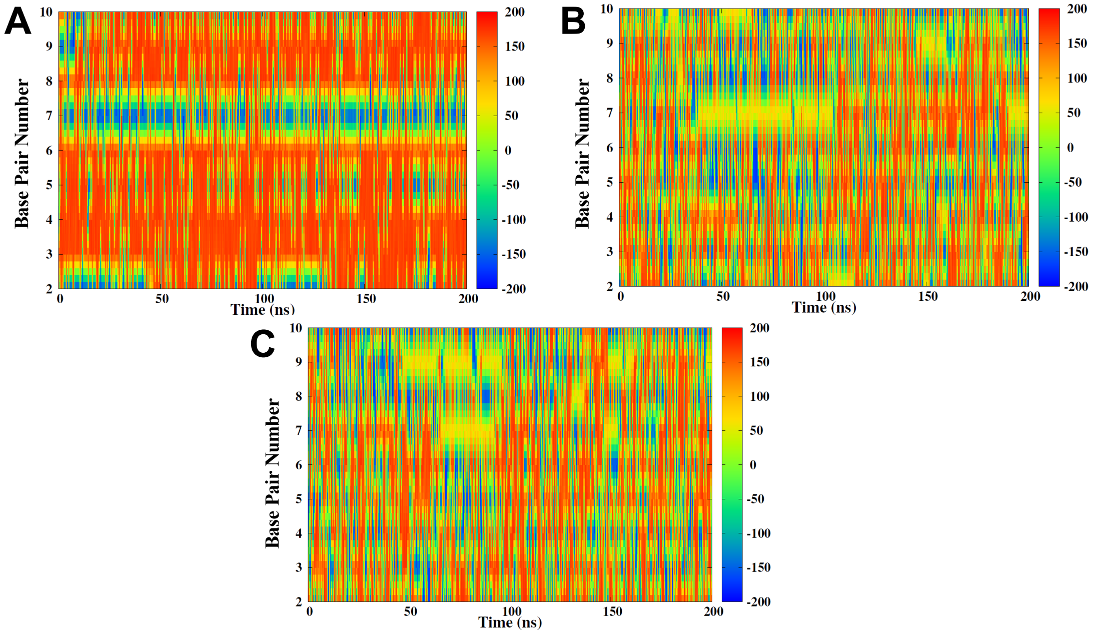

**Figure S3.** Time series of the backbone angle β for SASP/DNA (A), Naked DNA (B), and B-DNA (C).

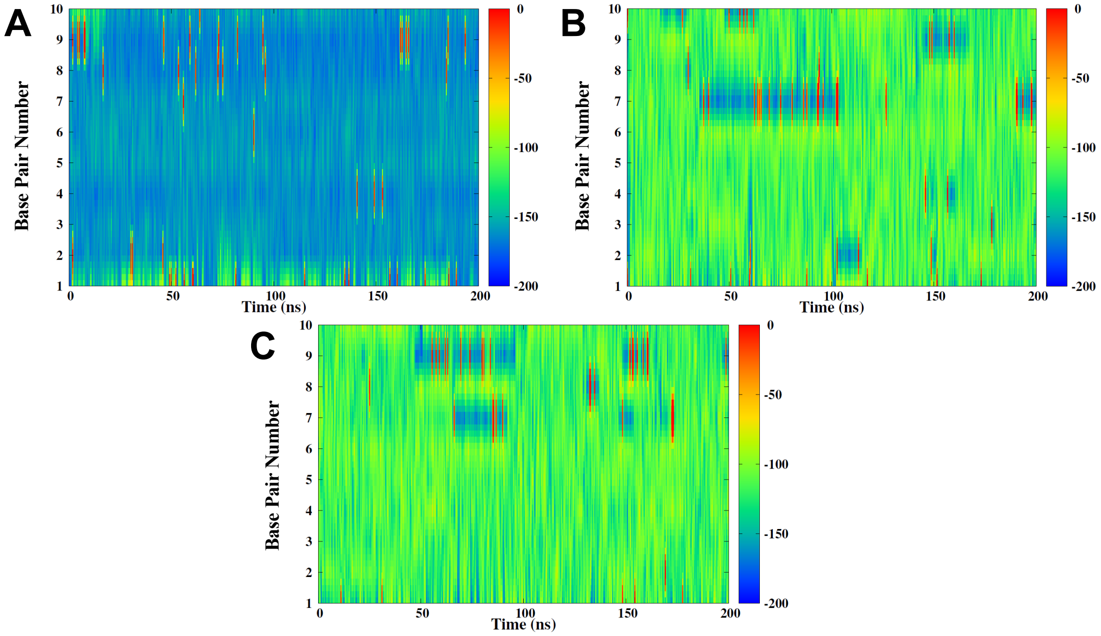

**Figure S4.** Time series of the backbone angle $\chi$ for SASP/DNA (A), Naked DNA (B), and B-DNA (C).

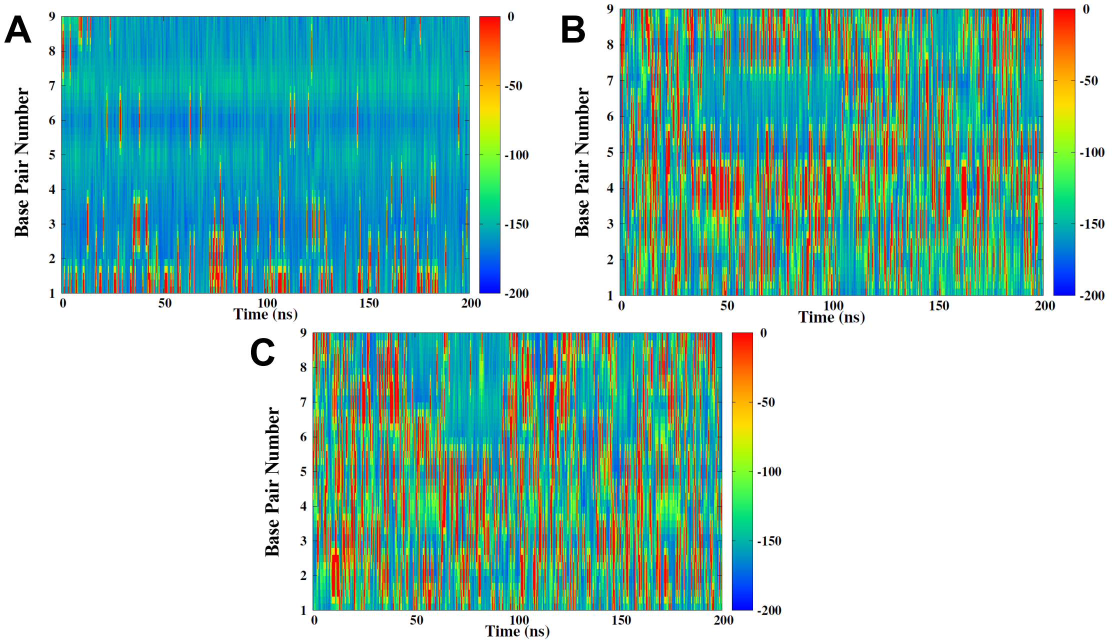

**Figure S5.** Time series of the backbone angle ε for SASP/DNA (A), Naked DNA (B), and B-DNA (C).

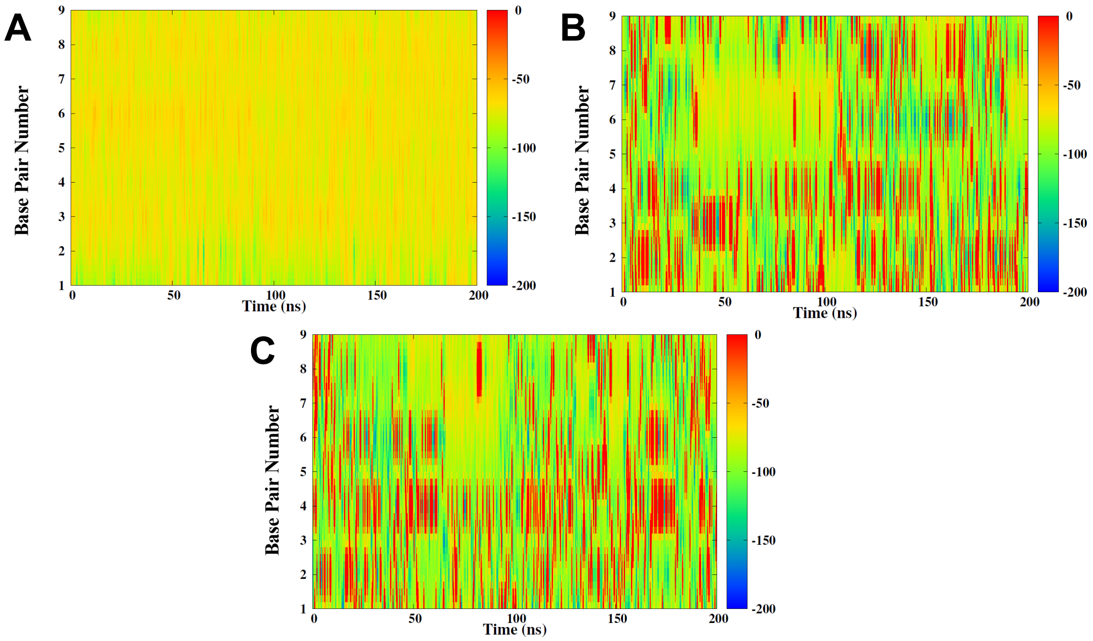

**Figure S6.** Time series of the backbone angle ζ for SASP/DNA (A), Naked DNA (B), and B-DNA (C).

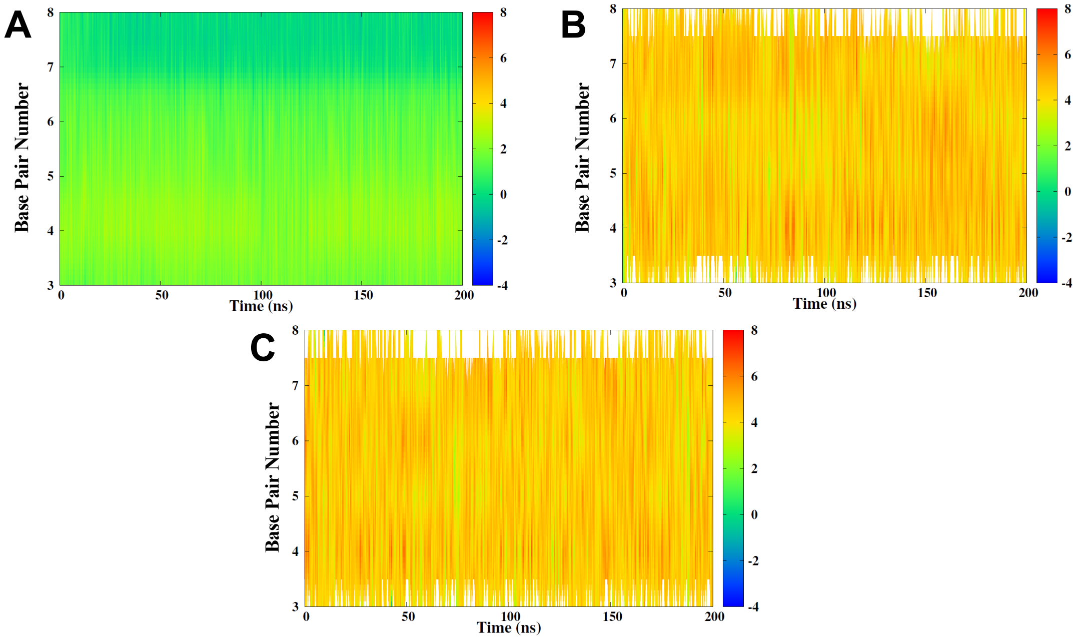

**Figure S7.** Time series of minor groove depth for SASP/DNA (A), Naked DNA (B), and B-DNA (C).

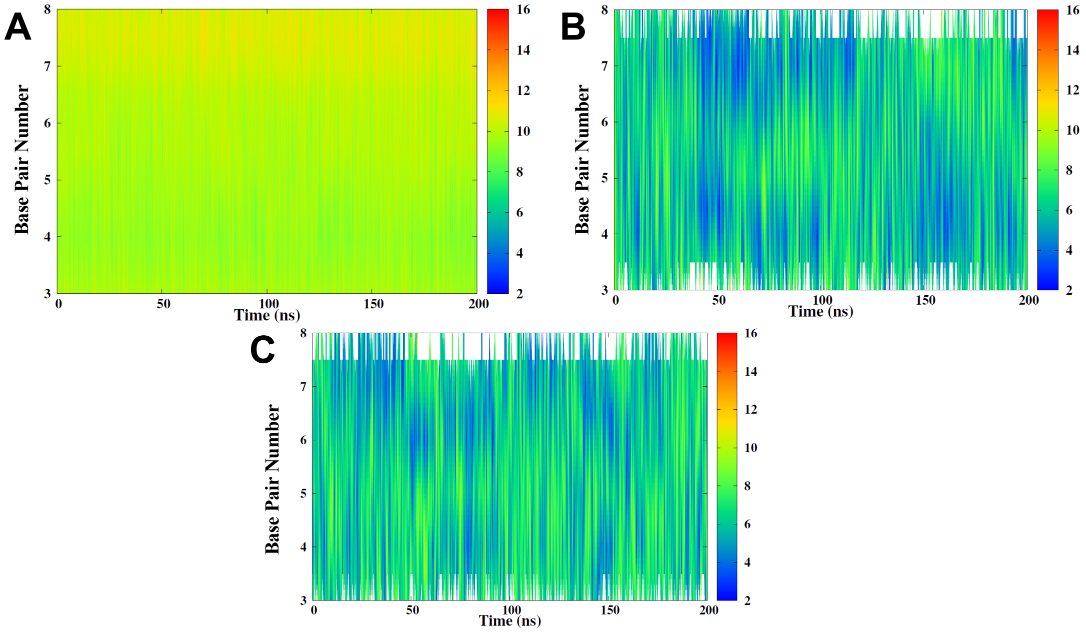

**Figure S8.** Time series of minor groove width for SASP/DNA (A), Naked DNA (B), and B-DNA (C).

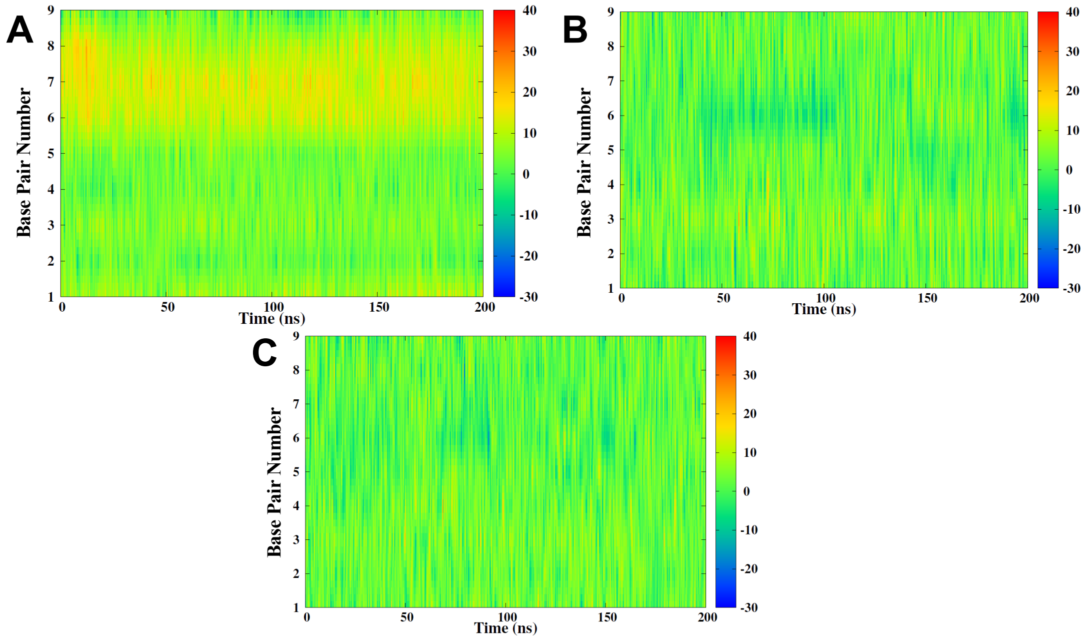

**Figure S9.** Time series of interbase roll for SASP/DNA (A), Naked DNA (B), and B-DNA (C).

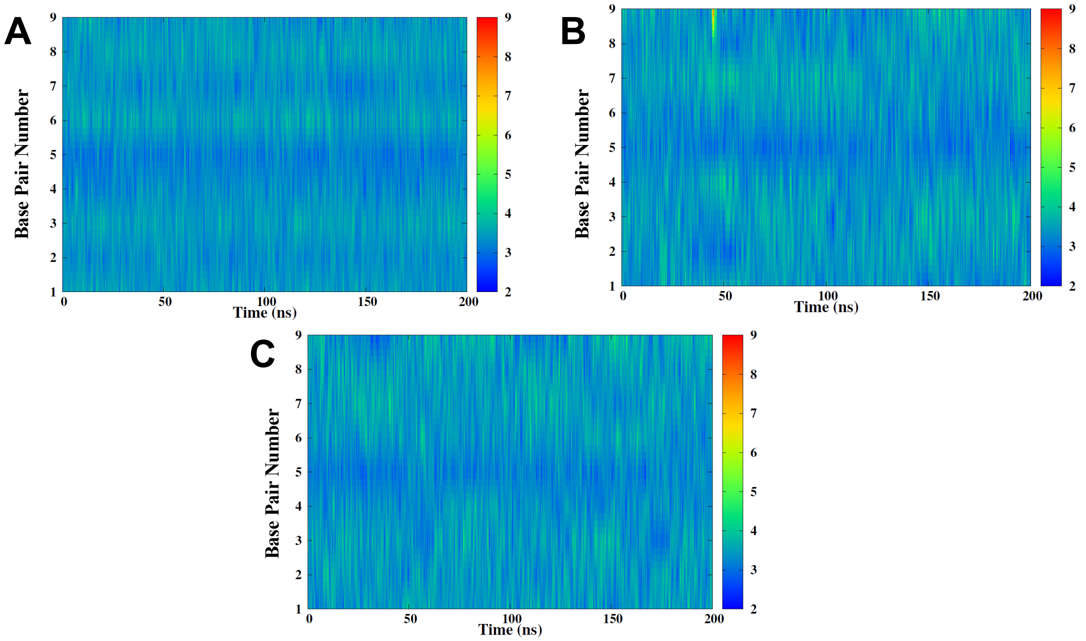

**Figure S10.** Time series of interbase rise for SASP/DNA (A), Naked DNA (B), and B-DNA (C).

#### Time series of nucleic acids' bending

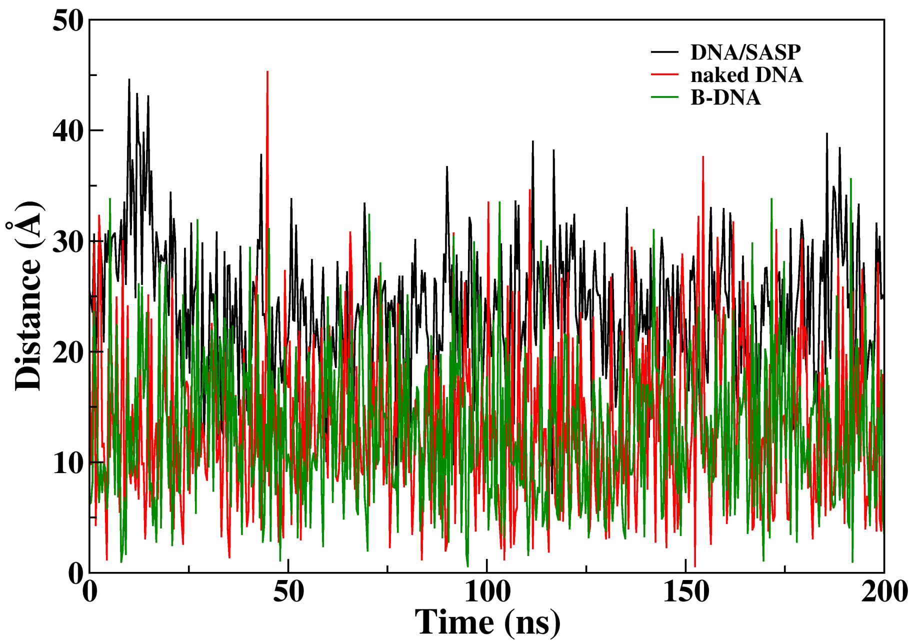

**Figure S11.** Time series of total bending for SASP/DNA (A), Naked DNA (B), and B-DNA (C).

#### Snapshots along the PMF for the B-DNA system

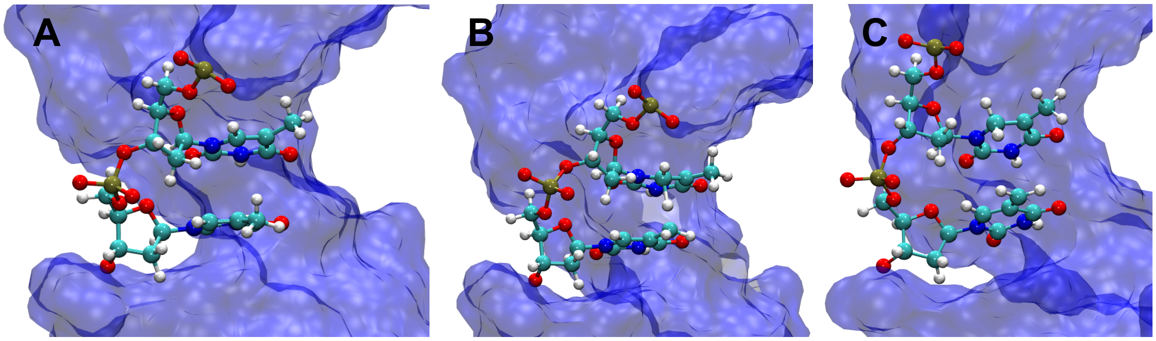

**Figure S12.** Representative snapshots extracted along the reaction coordinate for the B-DNA PMF representing the reactant (A), transition state (B), and product (C) regions.
